# Correcting for cell-type heterogeneity in epigenome-wide association studies: premature analyses and conclusions

**DOI:** 10.1101/121533

**Authors:** Shijie C Zheng, Stephan Beck, Andrew E. Jaffe, Devin C. Koestler, Kasper D. Hansen, Andres E. Houseman, Rafael A. Irizarry, Martin Widschwendter, Andrew E. Teschendorff

## Abstract

Recently, a study by Rahmani et al [1] claimed that a reference-free cell-type deconvolution method, called ReFACTor, leads to improved power and improved estimates of cell-type composition compared to competing reference-free and reference-based methods in the context of Epigenome-Wide Association Studies (EWAS). However, we identified many critical flaws (both conceptual and statistical in nature), which seriously question the validity of their claims. We outlined constructive criticism in a recent correspondence letter, Zheng et al [2]. The purpose of this letter is two-fold. First, to present additional analyses, which demonstrate that our original criticism is statistically sound. Second, to highlight additional serious concerns, which Rahmani et al have not yet addressed. In summary, we find that ReFACTor has not been demonstrated to outperform state-of-the-art reference-free methods such as SVA or RefFreeEWAS, nor state-of-the-art reference-based methods. Thus, the claim by Rahmani et al (a claim reiterated in their recent response letter [3]) that ReFACTor represents an advance over the state-of-the-art is not supported by an objective and rigorous statistical analysis of the data.

## Introduction

EWAS seek to identify DNA methylation (DNAm) changes which associate with a phenotype of interest, often encoded by a binary variable such as case-control status. The tissues used in such studies are heterogeneous mixtures of many underlying cell-types. This includes tissues such as whole blood [4], cord blood [5] buccal epithelium [6] and breast [7]. The proportions of these cell-types can vary between healthy individuals, as well as between cases and controls. The need for cell-type correction in these studies therefore arises due to the need for identifying differentially methylated CpGs (DMCs), which are not driven by changes in cell-type composition [4, 8].

Adjusting for cell-type heterogeneity can be performed using reference-based methods [9-12], which use a-priori defined reference DNAm profiles for the cell-types in the tissue of interest, or in a reference-free manner [13, 14]. Reference-based methods infer cell-type proportions in individual samples, which can be used as covariates in subsequent regression analyses to identify relevant DMCs. Reference-free methods on the other hand, infer from the data matrix “surrogate variables”, which capture variation due to cell-type composition, and which may also include variation due to other confounding factors. Reference-free methods don’t require pre-existing DNAm reference profiles and are therefore in principle applicable to any tissue type. They include tools such RefFreeEWAS [13], Surrogate Variable Analysis (SVA, ISVA) [15, 16], Removing Unwanted Variation (RUV) [17, 18] and more recently, ReFACTor [1].

All of these cell-type deconvolution methods make specific assumptions, which, if violated, render the method inapplicable. For instance, a reference-based method assumes that the main constituent cell-types of the tissue are known. The main assumption underlying the implementation of ReFACTor as presented by Rahmani et al [1], is that the top component of variation in the full data-matrix is driven by cell-type composition. It is a mathematical fact, that if the top component of variation is not correlated with cell-type composition, but with the phenotype or outcome of interest, that application of a method like ReFACTor will remove biological signal. We provided a clear example of this in the context of a breast cancer tissue EWAS [2], demonstrating that ReFACTor leads to a complete loss of power (sensitivity was estimated at less than 5%), in contrast to SVA which obtained a high sensitivity (~70-80%) to detect bona-fide breast cancer DMCs. This dramatic difference in performance can be easily understood, since SVA constructs surrogate variables (SVs) in the data variation space which is orthogonal to that associated with the phenotype of interest (here normal-cancer status). Thus, by construction, the SVs from SVA are unlikely to be associated with the phenotype of interest, and are much more likely to capture genuine confounding variation, such as that due to cell-type heterogeneity. Hence, application of ReFACTor is only possible on tissues and phenotypes where the top component of variation from a PCA is correlated with cell-type composition and not with the phenotype of interest. Although this problem could, in principle, be circumvented by applying ReFACTor to the controls only, this implementation of ReFACTor is not the one which Rahmani et al were originally presenting [1].

The original study by Rahmani et al assessed ReFACTor only in one tissue type: whole blood, a tissue for which the main assumption discussed above is likely to be valid in the context of most phenotypes. Indeed, the neutrophil fraction can vary substantially between individuals (as much as 35-40%, see e.g. Fig.6a in [12]), be they healthy or not. However, because the main constituent cell-types of whole blood are fairly well known and because high-quality reference DNAm profiles for these are available [19], this makes reference-based methods, such as the constrained projection (CP) technique of Houseman et al [9], the most attractive choice for EWAS conducted in whole blood. Rahmani et al performed a comparison of ReFACTor to Houseman’s CP algorithm in terms of their ability to estimate blood cell-type fractions, using only one dataset with matched DNAm and FACS-based cell counts, claiming improved modeling performance with ReFACTor. However, as we demonstrated in our letter [2], and as we argue again below, the assessment of ReFACTor in modelling cell-type composition is inherently biased because of overfitting. Thus, the claim by Rahmani et al that ReFACTor improves modeling of cell-type composition over a reference-based method is not supported by the data presented in their original letter, nor by the additional data presented in their response [3]. Indeed, as we demonstrate very clearly below, their recent analysis is as statistically and conceptually flawed as the one presented in the original letter.

## Discussion

### 1. ReFACTor breaks down in the cancer tissue EWAS setting

In an attempt to assess cell-type deconvolution algorithms in tissue types other than blood, we proposed to evaluate them in the context of a cancer tissue EWAS. Specifically, we constructed a list of gold-standard differentially methylated CpGs (DMCs) between normal breast and breast cancer cells, using cell-line data [2]. The need to use cell-lines arises because we require a comparison between samples that are as pure or clonal as possible and within a normal-cancer context. Thus, a comparison between normal mammary epithelial cells and breast cancer cell-lines represents the only viable approach to construct an approximate list of true DMCs between normal breast and breast cancer cells. Rahmani et al argue that our list of breast cancer DMCs is unreliable, because we compared 52 breast cancer cell-lines to only 2 normal controls.

We disagree that our list of breast cancer DMCs is unreliable. While we acknowledge that the use of 2 controls is not ideal, we nevertheless defined our list of gold-standard DMCs very carefully by demanding that differences in methylation between the two controls and the 52 breast cancer cell lines is large (at least a difference in mean methylation of over 50%). Indeed, the very fact that a method like SVA attained a sensitivity as high as 80% (as shown in Table 1 of our correspondence) in the independent breast cancer tissue set (consisting of 50 normals and 305 primary breast cancers) demonstrates that most of our gold-standard DMCs are bona-fide breast cancer DMCs. If they were false positives (as Rahmani et al claim), then these would not have validated with such high sensitivity in a completely independent dataset. In any case, all methods are being evaluated against the same set of gold-standard DMCs, so even if this gold-standard set contains a fraction of false positives, there is no reason to expect that this should drive marked differences in performance between methods (e.g. the 70 to 80% difference in sensitivity between SVA and ReFACTor). In other words, a number of false positives within our gold-standard list of true positives would only affect the absolute sensitivity, not the relative sensitivity between methods. The purpose of our Table-1 in Zheng et al [2] is therefore to demonstrate differences in the *relative* sensitivity between SVA and ReFACTor in the breast cancer tissue setting.

**Table-1:**
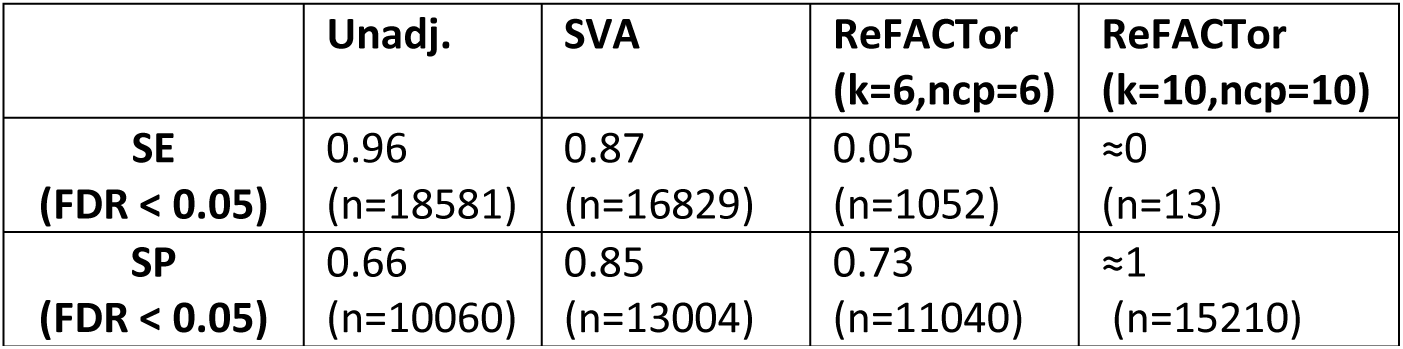
Table comparing the relative sensitivity (SE) and specificity (SP) of ReFACTor (for 2 different choices of k and ncp parameters, to SVA and to an unadjusted analysis. Sensitivity was estimated using a set of n=19379 true positives and 15212 true negatives, and are shown at an FDR < 0.05.

Second, Rahmani et al conduct a permutation analysis and conclude that many of our original gold-standard DMCs are false positives. However, it is expected that if we compare two breast cancer cell lines to a significant number of other breast cancer cell-lines that there will also be many DMCs. This follows from the known heterogeneity of breast cancer [20, 21]. In fact, any two breast cancer cell-lines will exhibit deviations from a normal state, and in general, some of the CpGs will be commonly altered and others will be unique to each cell-line. The key point to appreciate is that DMCs between 2 breast cancer cell-lines and a large number of other breast cancer lines will include loci which are indistinguishable between the 2 breast cancer cell-lines and the cell-lines representing a normal-state (in our case the mammary epithelial cell lines). In other words, these DMCs will themselves differ between normal and breast cancer generally, and therefore will validate when assessed in the independent breast cancer tissue EWAS, as observed by Rahmani et al [3]. Thus, in their permutation analysis, the resulting high sensitivity only reflects the fact that the DMCs derived from the permuted set constitute a different type of “true positive”. It follows that their permutation analysis does not refute in any way the reliability and quality of our original true positive DMC list.

In order to conclusively demonstrate this, we have used a slight extension of our previous approach to define a new gold-standard set of true positives. Instead of only using a list of cancer DMCs derived from a comparison of 2 normal to 52 breast cancer cell-lines (n=23,258) [2], we have intersected this list with a separate list of DMCs derived from the large breast cancer TCGA study [22] consisting of 81 normal samples and 652 breast cancers. This approach thus avoids the problem of only using 2 controls, it avoids the problem of cell-type composition and also avoids potential cell-culture artefacts. This strategy resulted in a new list of 19,379 true positive breast cancer DMCs, as well as a new list of 15,212 true negatives. We then used this new gold-standard list of breast cancer DMCs and computed the sensitivity of ReFACTor and SVA in our large breast cancer tissue EWAS study (50 normals and 305 primary breast cancers) [7] (Table 1). We obtained a sensitivity of 0.87 for SVA (similar to the value we estimated in Table 1 of our original correspondence [2]), whereas for ReFACTor we obtained a sensitivity value of only 0.04 (again in line with our original estimate [2]) (see also Table 1). Thus, this unequivocally demonstrates that the difference in power between the two methods is *not* due to us using only 2 control cell-lines, thus refuting Rahmani et al’s claim.

Most importantly, it is a mathematical fact that ReFACTor will remove biological signal, if this signal accounts for more data variation than cell-type composition. This is because ReFACTor effectively uses the top-PC to adjust the data for what it thinks is cell-type composition. As shown by us recently [7], variation in the adipose fraction only comes up in the 2^nd^ top-ranked PC, with the first component dominated by changes which reflect differences between normal and breast cancer epithelial cells (if the PCA is performed on normal breast samples only, then the top PC does correlate with adiposity). So, it is clear to us that ReFACTor would remove genuine biological signal in a cancer tissue EWAS setting, and this is likely to be the case for any cancer tissue EWAS, not just breast cancer.

Rahmani et al [3] further argue that our analysis in Table-1 is not meaningful because an unadjusted analysis performs optimally using a Bonferroni-adjusted threshold. However, it is critically important to realize that in the breast cancer-tissue EWAS setting we considered, that most of the data variation is due to the phenotype of interest (normal-cancer status) and not due to variation in cell-type composition. It follows that in an unadjusted analysis the top hits will be cancer DMCs. Thus, we would expect that a simple unadjusted analysis will probably be optimal under a very stringent Bonferroni threshold. This is because the top hits are not confounded by cell-type composition. The methods adjusting for cell-type composition would only add value over an unadjusted analysis if the thresholds are relaxed, because it is only upon relaxing the significance threshold that we would begin to capture DMCs which are affected by cell-type composition. Indeed, as shown in Table-1 of our correspondence letter [2], at a more relaxed FDR < 0.05 threshold, SVA’s specificity is over 20% larger than that of an unadjusted analysis, a result which we were able to confirm again using our new refined list of true negative CpGs (see Table 1), thus making it the preferred method. Thus, there is no paradox or contradiction, as claimed by Rahmani et al. Finally, we note that the use of a Bonferroni threshold is not recommended and not usual practice in a cancer *tissue* EWAS setting.

Rahmani et al [3] further perform an analysis, whereby they split a large set of breast cancer samples (n = 305) into two groups on the basis of the reference-based cell-composition estimates that we provided. One group was labeled as controls, and differential methylation effects were added to all samples in the other group in more than 20,000 sites. The authors then conclude that ReFACTor performs optimally in terms of specificity. However, this new analysis by Rahmani et al has three significant problems. First, it generates data for a scenario which is completely unrealistic, because as shown by us in Supplementary Figure 1 of our correspondence letter, the top component of variation in the breast cancer EWAS study does not correlate with variation in cell-type composition (adipose fraction). Adipose content is only correlated with the top-PC if the PCA is performed only on normal breast samples (i.e upon removing the breast cancers). Thus, Rahmani et al simulate an unrealistic scenario which is “tailored” for ReFACTor. Second, by comparing two random sets of breast cancers, there will be “true positive” DMCs, since breast cancers are highly heterogeneous. So, what the authors have done is to add another layer of “true positives” on top of the many biological ones that already exist, and it is therefore unclear to us which ones are more important for assessing sensitivity. It is also unclear which ones would be more highly ranked, thus rendering any sensitivity/specificity analysis completely futile and meaningless. Third, the manner in which Rahmani et al assess the cell-modeling composition issue is biased and incorrect, as we explained in our original letter, and as we explain again further below.

**Fig.1:**
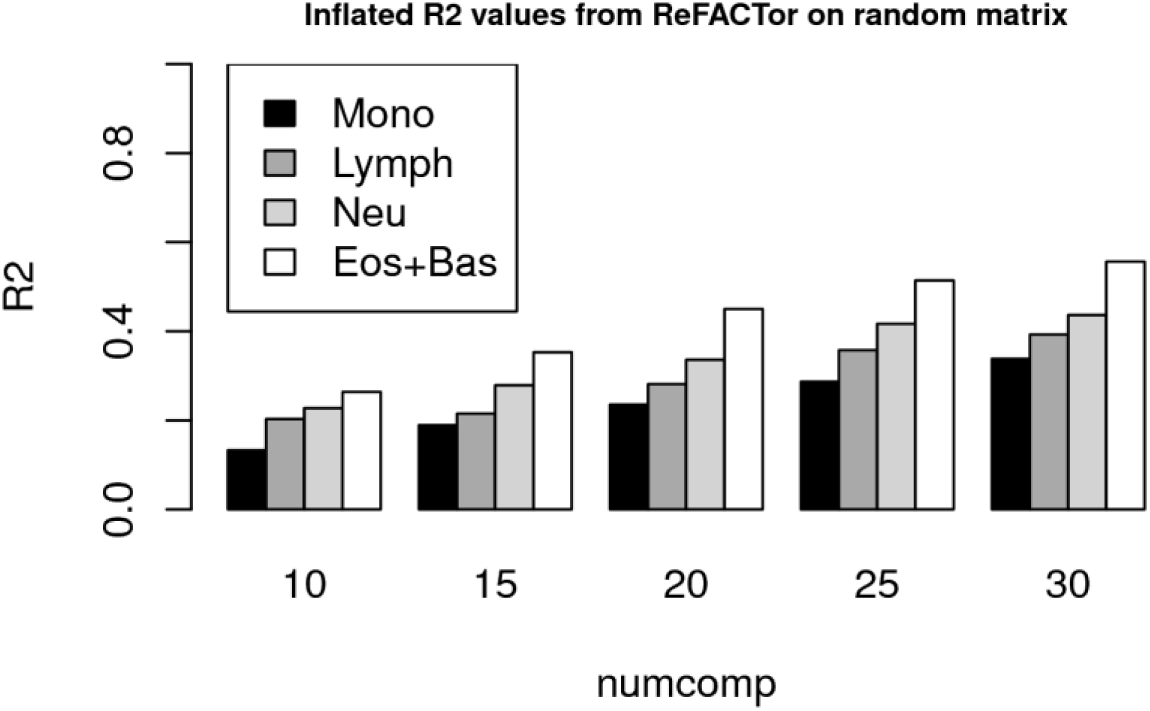
Inflated R^2^ values from ReFACTor on a random matrix. Barplot shows the estimated R^2^ values using ReFACTor and the procedure of Rahmani et al [1] on a randomly generated data matrix (uniformly distributed values on the interval (0,1)) of the same dimension as the GALA2 Infinium 450k set considered in Rahmani et al (number of CpGs ~470,000, number of samples =78). ReFACTor was run with the recommended parameters of k=6, and with numcomp varying from 10 to 30. This latter parameter determines the number of components used as predictors in the multivariate regression to model FACS cell counts, from which the estimated R^2^ values are obtained. Using the recommended value of numcomp=10, we see how R^2^ values obtained on a randomly generated matrix are inflated/biased by as much as 20%.

### 2. ReFACTor overfits and leads to inflated R^2^ values

When assessing cell-type composition and estimating R^2^ values against independent FACS cell counts, ReFACTor makes the following unjustified and unproven assumption, which leads to severely biased and inflated R^2^ values (as shown in Supplementary Figs.3 and 5 of our correspondence letter [2]). First of all, ReFACTor assumes that all the variation due to cell-type composition can be captured by only selecting the 500 CpGs which best fit the PCA decomposition of the data. This assumption, which is never proven, is the basis for then selecting an arbitrarily large number of ReFACTor components from the PCA performed on the reduced 500 CpG data matrix. Rahmani et al argue incorrectly that effectively all the components inferred from this PCA are linked to cell-type composition. We argue that the number of significant PC components needs to be determined using for instance a permutation based technique (as implemented for instance in SVA) [15] or RMT (Random Matrix Theory) [23]. Incidentally, the permutation approach, which derives an empirical null for the eigenvalue distribution of the data covariance matrix, and RMT, which uses an analytical null for the eigenvalue distribution, generally yield the same estimate for the number of significant PC components, as shown by us in Supplementary Figure 4 of our correspondence letter [2]. Alternatively, we proposed the LRT (Likelihood Ratio Test) approach to select relevant PCs, since this procedure avoids overfitting. Although no method is perfect, the key point is that by not selecting PCs, and instead postulating, unjustifiably so, that there are 10 or more biologically significant components correlating with cell-type composition (as ReFACTor does), ReFACTor suffers from severe overfitting, leading to substantially inflated R^2^ values. Indeed, the default parameter values used in ReFACTor [1] don’t even make biological sense: the original selection of 500 CpGs is based on how good they fit the full data based on the top 6 (k=6) PCA components, a number motivated by the fact that there are 6 main blood cell subtypes. However, later, in the 2^nd^ step, Rahmani et al perform a PCA on the 500 CpG reduced data matrix and extract 10 principal components (the “numcomp” parameter = 10), i.e a number larger than the original postulated number of blood cell-subtypes. The number of “10” was proposed without any statistical or biological justification. It is a statistical fact that adding even random components (which may well represent noise) to a multivariate model, will lead to inflated R^2^ values, because of overfitting. To make this point absolutely clear, we generated a random matrix of beta values between 0 and 1 and of the same dimension as the GALA2 Illumina 450k set (473838 CpGs and 78 samples) analysed by us previously [1, 2]. Applying ReFACTor to this random matrix with the parameter choices k=6 and numcomp=10, and correlating the resulting 10 “refactor” components to the GALA2 FACS cell counts for each of 4 blood cell subtypes, resulted in R^2^ values of 0.13, 0.20, 0.23 and 0.26 for Monocytes, Lymphocytes, Neutrophils and Eosinophils, respectively. These inflated R^2^ values increase in line with the number of components included in the model (Fig.1).

This demonstrates the importance of using a technique like RMT to estimate the number of significant components. Indeed, using RMT on the generated random matrix (i.e. all values identically and independently distributed) we obtain zero significant components, which leads to the correct conclusion that R^2^=0 for an artificially generated random data matrix which should show no association with cell-count estimates. We further note that the inflated R^2^ values of 10-20% obtained here are consistent with those we estimated previously (Supplementary Figure 3 in our correspondence letter shows that ReFACTor leads to inflated R^2^ values of 20% or higher), which was confirmed further by using an unbiased training-test set strategy (Supplementary Figure 5 in our correspondence letter).

Rahmani next argue that they have redone the analysis using a larger set of 560 samples. However, what they don’t disclose in the main letter is that true FACS cell counts are only available for the same 78 samples used in their original letter, and that for the rest of the samples, cell counts were imputed, which once again does not allow for an objective comparison. Moreover, although we agree that by using more samples we are likely to pick out more significant components, the problem remains that these need to be selected using a criterion such as LRTs or permutations/RMT. Rahmani et al do not do this and therefore their subsequent assessment is once again severely overfitted, leading to an apparently high R^2^ value and improved performance over Houseman’s algorithm [9].

Rahmani et al next argue that the number of dimensions estimated by RMT is linearly determined by sample size, making it inapplicable. In response to this, we have simulated two datasets with two components of variation each, and which differ in terms of sample size (n=20 & n=200 samples). In both cases, Random Matrix Theory (RMT) [23] correctly predicts 2 components (Fig.2), providing a clear counterexample to the statement by Rahmani et al:

**Fig.2:**
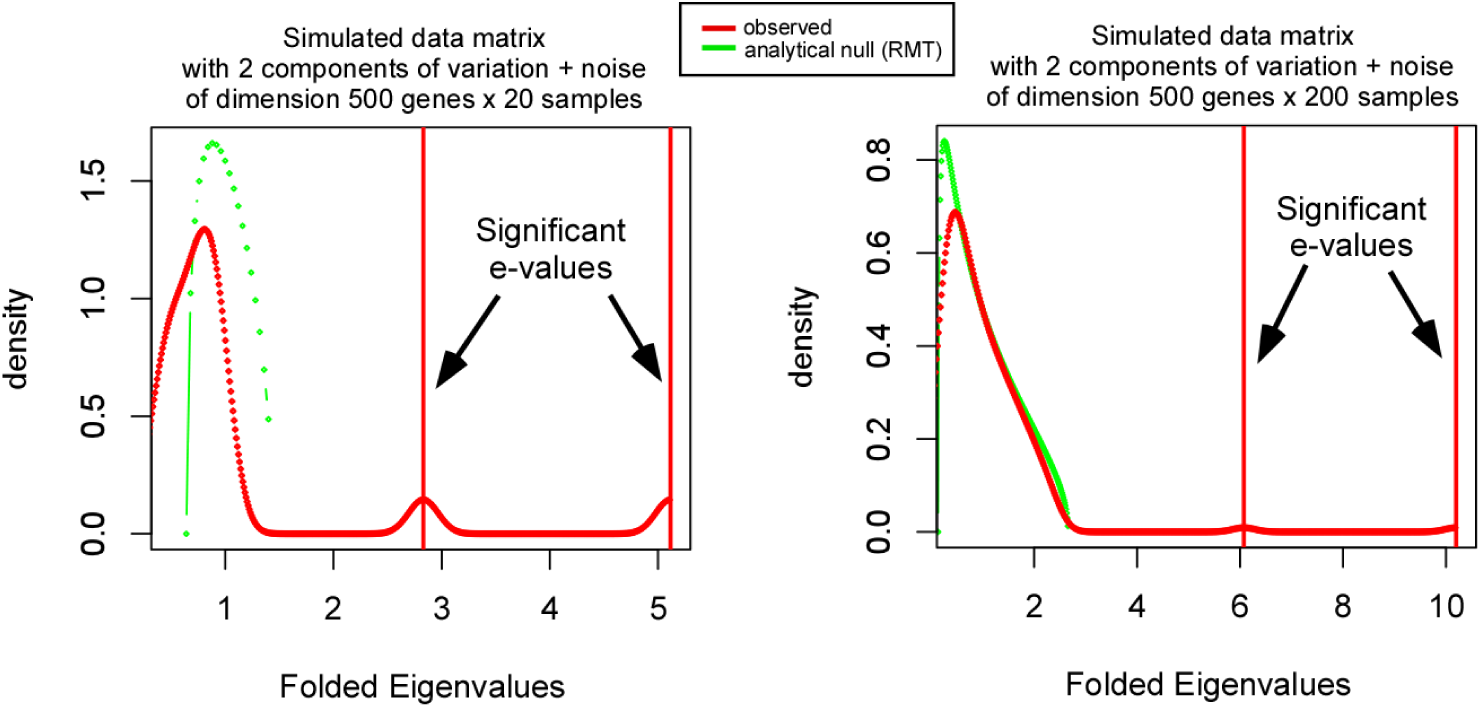
RMT estimates of the number of significant components of variation in two artificial datasets, each with two simulated components of variation, but which differ in the number of samples (20 & 200), as indicated. Green points and curve indicate the null eigenvalue distribution, as predicted by RMT. The red data points and curve is the eigenvalue distribution of the observed data covariance matrix. As can be seen, in each case there are two peaks with values larger than the maximum expected for a random matrix (green line). Thus, in both cases RMT predicts 2 significant components of variation.

Moreover, the claim by Rahmani et al that the RMT dimensionality estimate increases linearly *every time a sample* is added is untrue. Indeed, the plots shown in Supplementary Figure 6 of Rahmani et al’s response [3], clearly demonstrate that the dimensionality increases in a step-wise fashion, i.e. *the dimensionality does not increase every time 1 sample is added*. Second, all datasets considered in their figure are very large datasets, for which the expected number of components should also be large. Interestingly, the authors choose to stop assessing the RMT estimates well before the maximum sample size of the study. For instance, in Liu et al [4] there are 689 samples, yet Rahmani et al only plot the RMT estimates up to about n=150. <u>The dimensionality of a dataset is expected to increase with sample size in line with the fact that there will be additional chip effects, batch effects and genetic effects which become more prominent as the number of samples is increased</u>. Indeed, for the large dataset of Liu et al (~480,000 CpGs × 689 samples), RMT estimates 60 significant components. Given that with Illumina 450k data we would need at least 58 separate beadchips (12 samples per 450k beadchip), our estimate of 60 significant PCs is actually strongly consistent with a large number of chip effects. Once we factor in biological sources of variation and other technical artefacts, the RMT estimate of 60 significant components makes a lot of sense. To confirm this, we correlated each of the 60 PCs to biological and technical factors, which unequivocally demonstrates that even low ranked components (e.g. PC-58) still correlate very strongly with beadchip effects (Fig.3).

**Fig.3:**
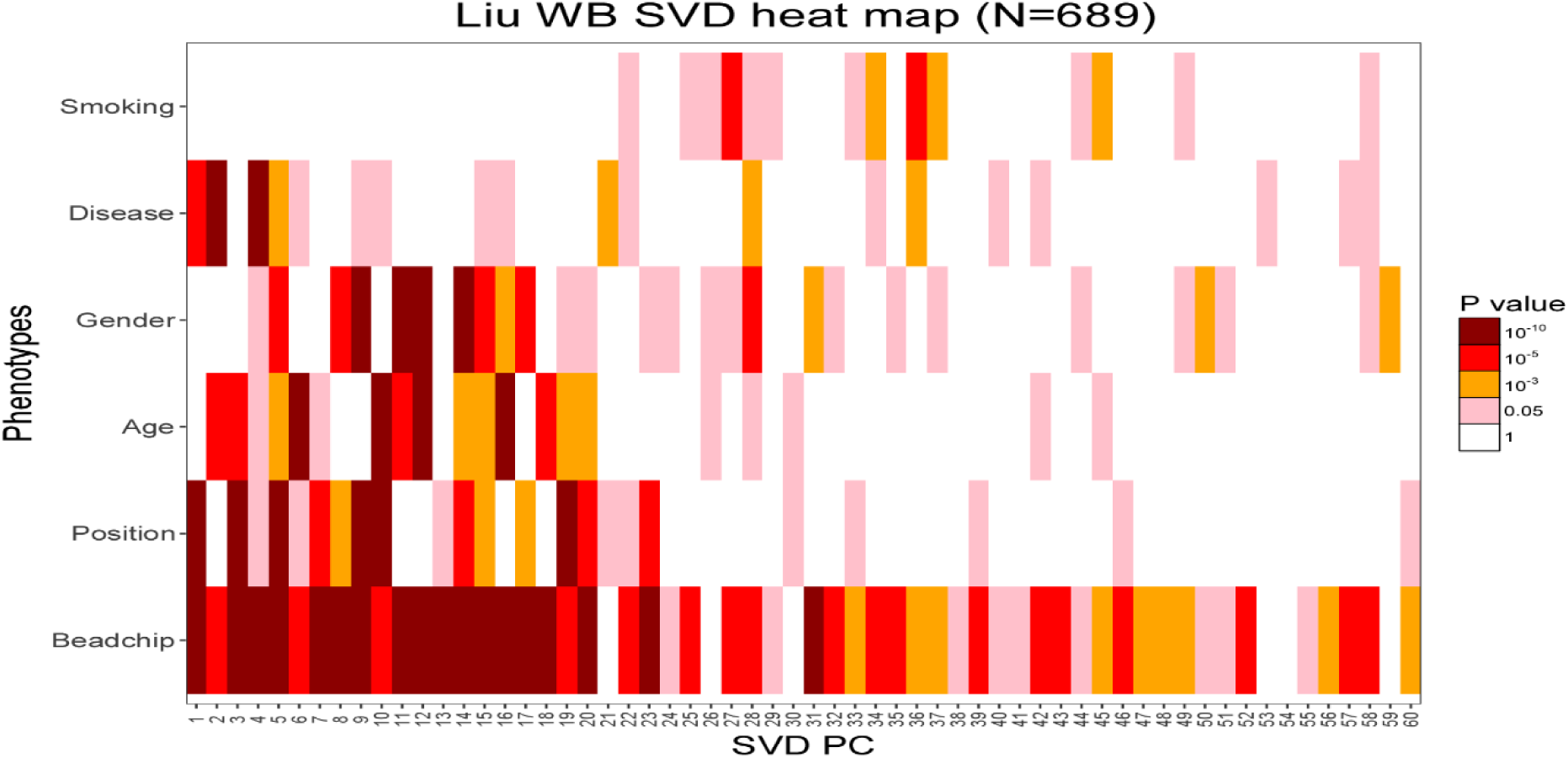
Heatmap of P-value associations between significant principal components from a PCA/SVD on the large 450k data set of Liu et al [4] against a number of different biological and technical factors. PCA/SVD was applied over all ~480,000 CpGs and 689 samples. RMT estimated a total of 60 significant components and observe how components ranked as low as 58 still correlate very strongly with beachip (P<1e-5).

Supporting this further, other large studies performed in whole blood also demonstrate large numbers of significant components of variation which often capture genetic variation [24]. So, in our view, the observed linear increase in dimensionality with sample size in these large *real* datasets should be expected and is correct.

A potential valid criticism against RMT could be that it compares the observed eigenvalue distribution of a PCA against an analytical null distribution which is derived in the limit of large matrices [23]. Hence, for “small” matrices the analytical null could lead to biased estimates. However, what constitutes “small” needs to be determined, not assumed. An alternative approach to RMT which does not depend on the size of the data matrix is to scramble up the data matrix (the Buja-Eyuboglu (BE) algorithm used by Leek JT et al [15]), and compare the resulting eigenvalue distribution to that of the original data matrix (using many randomizations to derive an empirical null eigenvalue distribution). As shown in Supplementary Figure 4 of our letter, the RMT and BE-based estimates agree perfectly, even on datasets of relatively small sample size (n=6 & n=12), thus justifying the applicability and validity of RMT on these “small” data matrices.

**Fig.4:**
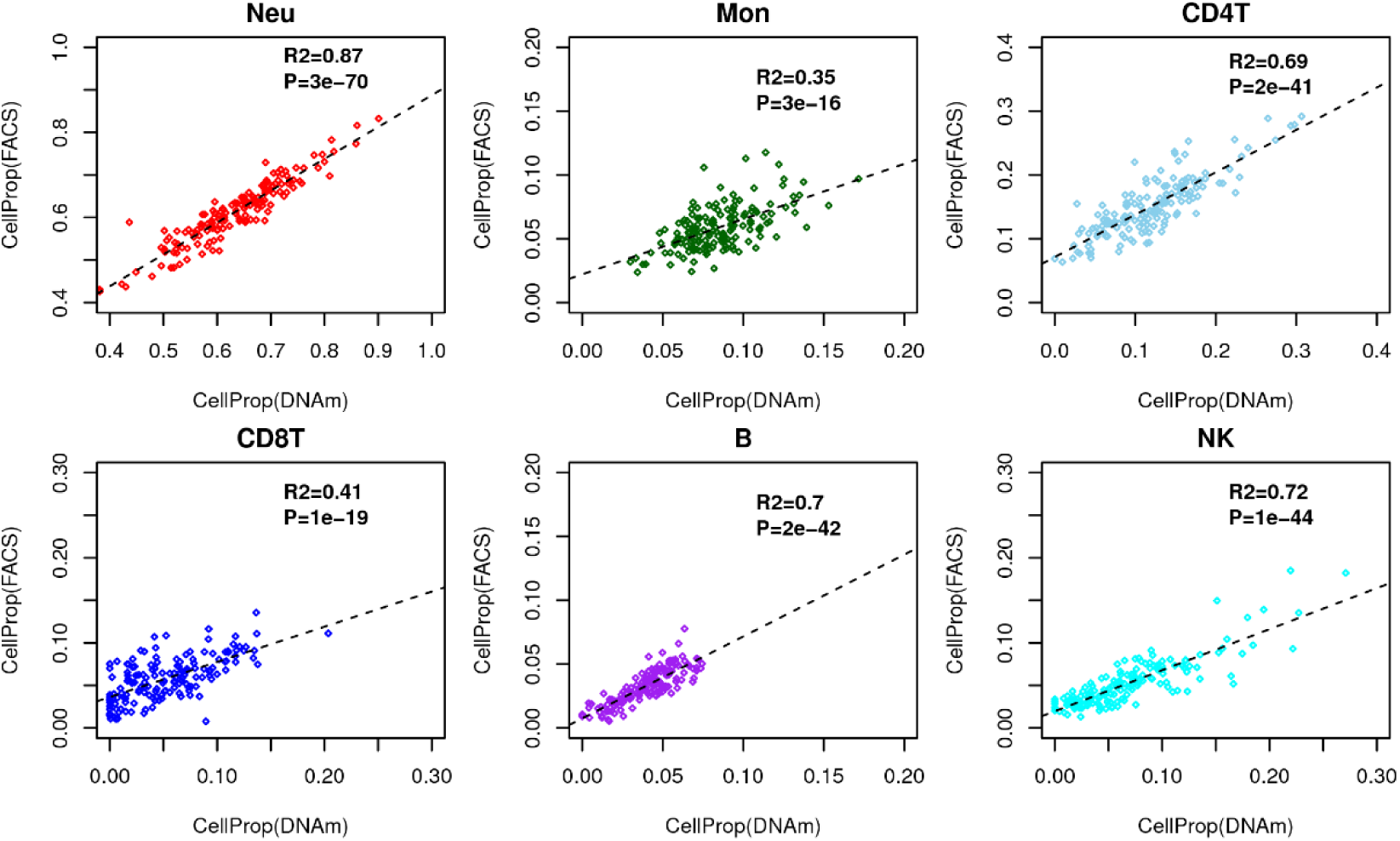
Scatterplots of blood cell subtype fractions as estimated with FACS (y-axis) against the corresponding estimates obtained using Houseman’s reference-based method, using the same blood cell subtype reference-database as used by us previously [2, 12] in a whole blood Illumina EPIC dataset of 162 samples. R^2^ values and linear regression P-values are given. This figure demonstrates that high R^2^ values are achievable for multiple cell-types in a dataset which is larger than the GALA2 set, thus raising questions to the reliability of the cell counts in the GALA2 set.

### 3. Inappropriate comparison to a reference-based method

Next, Rahmani et al used five large whole-blood data sets and divided the samples in each data set into two groups on the basis of cell-composition distribution [3]. Then, they conducted an EWAS on the assignment into groups as the phenotype. In this scenario, the assignment into groups is expected to be correlated with the true underlying cell composition, and an insufficient correction will lead to spurious associations. They conclude that ReFACTor consistently outperformed Houseman’s (CP) reference-based method; particularly, the reference-based method resulted in more than 1,000 false positives in some cases, whereas ReFACTor resulted in a few dozen at most. This specific analysis performed by Rahmani et al once again considers a scenario for which ReFACTor was designed, therefore, it is unsurprising that it performs well. Indeed, the scenario considered here is one where (i) the top component of variation is associated with cell-type composition (by construction and design), and (ii) where there might be additional batch effects (since the data is drawn from real datasets). In this specific scenario, as acknowledged by us in our letter [2] and recent review [12], ReFACTor will do reasonably well in terms of sensitivity and specificity. In fact, *Houseman’s CP method may not outperform ReFACTor, because it would not correct for additional batch effects*. The fair comparison would have been to assess ReFACTor against the combined application of a reference-based method plus a batch-correction algorithm such as COMBAT [25]. However, Rahmani et al chose not to present a fair comparison.

### 4. Low R^2^ values in GALA2 dataset

The data from Koestler et al [26] contains matched FACS cell counts and DNAm data. In this set, we observed relatively high R^2^ values (typically around 0.8) for most of the blood cell subtypes (Supplementary Figure 8 in our letter [2]), whereas in the GALA2 dataset R^2^ values are much lower hovering around 0.4 [2]. Had we observed this difference in only one cell-type, we would agree with Rahmani et al that a difference in R^2^ values of 0.4 could be a statistical fluke due to the small sample size of the Koestler set (n=6). However, this difference is observed across several cell-types, thus raising the question about the quality of the FACS cell counts in the GALA2 dataset. Indeed, Rahmani et al acknowledge that the cell counts and DNAm profiles for the 78 samples from GALA2 set were performed many months apart, during which time cell-counts could have varied substantially. Indeed, blood cell-counts are known to vary as a function of season, the time in the day in which the sample was drawn, infections, medications, allergic reactions, exercise and stress (see e.g. [27]). To confirm that the FACS cell counts of the GALA2 dataset are potentially of low quality, we recently obtained very high R^2^ values (>0.7) for 4 different blood cell subtypes in an independent study profiling a larger number of whole blood samples (n=162) with matched FACS cell count data (Fig.4). Based on this, we seriously question the suitability of the GALA2 cell counts as a “gold-standard” and for drawing any reliable conclusions.

### 5. Inappropriate choice of gold-standards in real EWAS

In the original letter, ReFACTor was further assessed using an inappropriate choice of gold-standard DMPs in real data. As shown in Figure 2 of Rahmani et al [1], the authors used an EWAS of Rheumatoid Arthritis (RA) [4]. However, this analysis is not conclusive, as there is no gold-standard set of CpGs that have been unequivocally demonstrated to be associated with RA. In fact, in this analysis the authors resort to using the reference-based (CP) result of Liu et al [4] to “validate” ReFACTor. Thus, this analysis can’t be used to support the claim that ReFACTor outperforms the reference-based method, nor that it outperforms any of the other reference-free methods. A better approach would have been to focus on smoking, for which there are by now at least 5 to 6 independent whole blood EWAS all leading to the same set of smoking-associated CpGs [28]. Although we acknowledge that a fraction of these consistent smoking-DMCs may be caused by shifts in blood cell-type composition not accounted for by current blood reference DNAm databases, this is still preferable over using a list of CpGs which have not yet been reproduced by an independent study. In fact, as shown by us [6], a large fraction of smoking-associated DMCs do not map to genes that have been reported to be markers for blood cell subtypes, so it seems likely that a considerable fraction of smoking-DMCs are not the result of shifts in blood cell subtypes. Focusing on smoking-associated DMCs would in our opinion provide a more objective “gold-standard” list for assessing methods in studies where detection of these sites is compromised due to intra-sample cellular heterogeneity and other confounding factors.

### 6. No comparison to state-of-the-art methods

The study by Rahmani et al claimed improvement over the state-of-the-art, a statement which is reiterated in their recent response letter. However, the fact remains that ReFACTor has not been compared to other conceptually similar algorithms such as RUV [18], of which there are a number of different variants [17]. Recently it was noted that RUV can empirically estimate and control for cell-type composition [8]. One version of the RUV method [8] infers “control” probes that are affected by the confounder (cell composition) but not the outcome of interest. Thus, this is conceptually very similar to ReFACTor, and that no direct comparison to RUV was ever offered, further calls into question the conceptual advance of ReFACTor. Likewise, in the original letter, ReFACTor was not compared to SVA [15], a state-of-the-art method, which in the meantime has been found by an independent study to be the most robust method [29], in line with our recommendations.

## Conclusions

As elaborated in the Supplementary Information document of our letter [2], the choice of a cell-type deconvolution method depends mainly on 3 factors: (i) tissue-type, (ii) phenotype of interest and (iii) the information sought by the study. The first two factors determine the relative data variation between phenotype (e.g. cancer/normal status, smoking exposure, type-2 diabetes) and cell-type composition. The third factor refers to whether cell-type composition estimates (absolute or relative) are desired in addition to the identification of DMCs.

In scenarios where cell-type composition drives most of the variation and where the underlying main cell subtypes are known, a reference-based method is, in our view, the current state-of-the-art since this approach should yield fairly reliable cell-type composition estimates, as well as a reliable set of DMCs. This recommendation is further supported by a recent letter by Hattab et al [30]. We further note that a reference-based method can be easily combined with batch-correction methods such as COMBAT if evidence for batch effects is present. If however there is evidence of additional, potentially unknown, confounding factors, then an approach such as SVA or RUV should be preferable for the identification of DMCs, a view also supported by a recent comparative study [29]. Algorithms such as SVA should also be preferable in scenarios where a reference DNAm database is not available and where it is unknown how much relative variation is associated with phenotype and cell-type composition.

In summary, ReFACTor is not “state-of-the-art” and in our opinion only applicable in the specific case where (i) estimates of cell-type fractions are not needed, (ii) where it is known with <u>certainty</u> that the largest amount of data variance is associated with cell-type composition and not with the phenotype of interest, and (iii) if there is evidence for additional unknown confounding factors. However, this also constitutes a scenario where tools such as SVA or RefFreeEWAS have been shown to perform equally well.

